# Effects of goldenrod (*Solidago gigantea* Aiton and S. *canadensis* L.) invasion on the wet meadow vegetation composition of Somogyfajsz wood pasture

**DOI:** 10.1101/2024.05.10.593301

**Authors:** Katalin Szitár, Melinda Kabai, Zita Zimmermann, Gábor Szabó, Bruna Paolinelli Reis, László Somay

## Abstract

In recent decades, restoring abandoned wood pastures are being rediscovered as an effective way to preserve biodiversity. Their restoration include the control of invasive plant species. In our study, we compared the vegetation composition of wet meadow patches uninvaded and invaded by giant and Canadian goldenrod in a wood pasture located in Somogyfajsz, Hungary. We sampled the vegetation of the wet meadows of the pasture in 16 pairs of 1 x 1 m invaded and uninvaded quadrats in June 2020. We studied the effect of goldenrod invasion on nature conservation value in terms of species diversity, origin of species, and Social Behaviour Types, and forage quality and quantity. The study revealed that goldenrod invasion had negative effects on species richness and community-level biomass in wet meadows, leading to a significant reduction in species diversity and total cover. Goldenrod invasion also impacted the richness and cover of both native and exotic species, with a decrease in richness but not cover, highlighting the competitive ability of goldenrods and their potential to alter vegetation composition.

## 1 INTRODUCTION

Wood pastures represent an essential component of cultural and ecological legacy in Europe (Bergmeier et al., 2010). They are tree-lands with a diverse forms ranging from natural grasslands with scattered trees and tree groups to closed-canopy woodland (Garbarino & Bergmeier, 2014). They can be characterised by high habitat, ecotone and structural heterogeneity, which results in exceptionally high species diversity. They are silvopastoral systems traditionally managed with parallel grazing and forestry practices (Hartel et al., 2015). In Central Europe, this land use was common practice until the 19^th^ century when it was banned due to growing timber and wood demand (Bergmeier et al., 2010). This coupled with a decline of livestock farming resulted in cessation of grazing and forest expansion on wood pastures (Garbarino & Bergmeier, 2014). In recent decades, however, wood pastures are being rediscovered as an effective conservation intervention to preserve biodiversity and deliver various ecosystem services (Plieninger et al., 2015). Restoration of wood pastures include the reintroduction of traditional management practices such as grazing with traditional livestock breeds, and control of invasive plant species.

Biological invasion is one of the most important drivers of biodiversity loss worldwide (IPBES, 2019). Giant goldenrod (*Solidago gigantea* Aiton) and Canadian goldenrod *(Solidago canadensis* L.) are clonal perennial herbs of North American origin that have invaded large areas in Europe but have also naturalized in Japan, Korea, Russia, Australia, and New Zealand (Weber & Jakobs, 2005). The species was introduced to Europe in the 18^th^ century as an ornamental species (Szymura et al., 2015). Giant goldenrod is a typical wetland species in its native range invading also dry grasslands besides wet meadows in the invaded range (Weber & Jakobs, 2005). It occupies a wide range of habitats from nutrient poor to nutrient rich but its presence is concentrated on nutrient rich wet soils (Scharfy et al., 2009).

Both goldenrod species are considered a pest species with serious environmental and economic impact on grasslands where it forms large and monodominant stands (Botta-Dukát, 2016; Kundel et al., 2014). Earlier studies showed that goldenrod invasion resulted in the decrease of plant diversity in old-fields and altering successional trajectories (Fenesi et al., 2015, Gusev, 2015). Vanderhoeven et al. (2006) showed that giant goldenrod invaded stands had 2-3-fold higher aboveground productivity and higher nutrient pools in standing phytomass than uninvaded stands. However, he also noted that it affected topsoil chemistry only modestly. Similarly, Canadian goldenrod had only a limited effect on soil seed bank in either wet meadows or anthropogenic habitats according to Kundel et al. (2014).

In this study, we compared the vegetation composition of wet meadow patches uninvaded and invaded by giant and Canadian goldenrod in a wood pasture located in Somogyfajsz, Hungary. We specifically hypothesised that goldenrod invasion resulted in the decline in nature conservation value in terms of species diversity, origin of species, and Social Behaviour Types sensu Borhidi (1995), and forage quality and quantity.

## 2 MATERIALS AND METHODS

Our study was located in the Somogyfajsz wood pasture in the Pannonian biogeographical region in the Inner Somogy Region, Southwest Hungary (WGS 46°29’N, 17°33’E). The pasture is part of the Hungarian Ecological Network, and the Natura 2000 network under the Birds Directive (Special Protection Area code HUDD10008). The climate is continental with a sub-Mediterranean character, with mean annual temperature of 10.6 ºC, and mean annual precipitation of 700-720 mm (Kovács-Láng, 1980). The soil of the pasture is blown acidic sand (Arenosol) with low humus content (below 1%). The semi-natural vegetation of the landscape is composed of oak-hornbeam deciduous forests (*Fraxino pannonicae-Carpinetum*) developed on rust brown forest soil, and acidic sand grasslands (*Festuco vaginatae-Corynephoretum*) developed on sand hilltops and sides (Kovács-Láng, 1980). The water-impacted areas are covered with *Alopecureto-Festucetum* wet meadows and black alder swamps (*Carici elongatae-Alnetum*; Juhász and Dénes, 2006).

The vegetation of the wood pasture was a mosaic of alder forests and wet meadows in the 18th century, and a grazed meadow during the 19th century according to the Habsburg Military Survey maps (Biszak et al., 2014). The rehabilitation of the wood pasture began in 1993 by clearing wet meadows invaded by shrubs and alders to recreate grazing area. The area is grazed extensively by a herd of approx. 80 Hungarian Grey cattle, and is mowed usually once a year late summer. The sampling site is a five hectares’ wet meadow dominated by *Festuca pratensis, Deschampsia caespitosa, Agrostis stolonifera*, and *Carex* spp. Giant and Canadian goldenrod invasion has started at least 20 years ago, and resulted in scattered goldenrod patches within the area.

We sampled the vegetation of the wet meadow in 16 pairs of 1 x 1 m relevés in June 2020. The pairs were designated at edges of goldenrod patches. One of each pair (invaded) was located within the goldenrod clone with goldenrod cover at least 50 %. The other pair was designated as close to the other one as possible without any goldenrod shoots (uninvaded). We made coenological relevés in each plot with an estimation of aboveground percentage cover of each vascular plant species, the cover of bare soil and plant litter. We also sampled the biomass of the vegetation next to each relevé in three 15 x 30 cm quadrats. We clipped the vegetation at the height of 1 cm aboveground. We sorted the clipped material into three fractions: goldenrod, other herbs, and graminoids (*Poaceae* and *Cyperaceae*). We weighed the fractions with an accuracy of 0.01 g after drying the fractions at 65 ºC for 48 h.

We grouped the species based on several species characteristics related ecological condition and pasture quality: origin (native vs exotic), life span (short-lived vs perennial), growth form (graminoid, legume, herb), and social behaviour types (hereafter SBT) sensu Borhidi (1995). The SBT categories reflect the species’ ecological role in the vegetation. We applied the following SBT categories: competitors (dominant species in natural communities), specialists (stress tolerants with narrow tolerance and low competitiveness), generalists (stress tolerants with wide tolerance), and ruderals. The latter including natural pioneers (associated with initial successional stages after natural disturbances), disturbance tolerants (plants of natural habitats with anthropogenic disturbance), weeds (native anthropogenic species), ruderal competitors (native community-forming dominant weeds), and alien competitors (habitat transforming invasive species). We collected species traits from the PADAPT database (Sonkoly et al., 2023), the Flora database 1.2. (Horváth et al., 1995), and the identification guide of the Hungarian flora (Király, 2009). We calculated total species richness, the number and relative cover of each functional group categories for each relevés. Goldenrod was not assigned to any trait groups, total species richness or cover in the analyses.

We tested the effect of goldenrod presence on total species richness, and the richness of species assigned to each trait value using general linear mixed effects models (glmer) with Poisson distribution and log link function using the lme4 package of Bates et al. (2015). Potential overdispersion of the errors was tested with the DHARMa package (Hartig, 2022). We also tested for the effect of goldenrod presence based on total species cover, percentage cover of each trait value, and herb, and graminoid biomass. For these analyses, we used linear mixed effects model (lme) in the nlme package (Pinheiro et al., 2012). The model residuals were visually checked for homoscedasticity, and in the case of biomass and cover data, for normality of errors. The analyses were conducted in the R programming environment (R Core Team, 2022).

## 3 RESULTS AND DISCUSSION

The average goldenrod cover of the invaded relevés was 75.5 +/- 2.6 (SEM). We found negative effects of goldenrod invasion on species richness and community-level biomass but only a single effect on aboveground plant cover of wet meadows in the wood pasture. We observed altogether 85 vascular plant species in the 32 1 x 1 m relevés, of which 20 species occurred exclusively in uninvaded, whereas 16 species only in invaded plots (Table S1). Despite the similar size of the species pool, we found that goldenrod invasion reduced average species richness by 32.5% in the 1 x 1 m plots (Table 1). Similarly, Fenesi et al. (2015) found a negative correlation between the density of goldenrods and the number of plant species richness in Romanian old-fields irrespective of their successional age. De Groot et al. (2007) also showed that plant species richness were significantly lower under dense goldenrod stands than in undisturbed semi-natural vegetation.

**TABLE 1.**
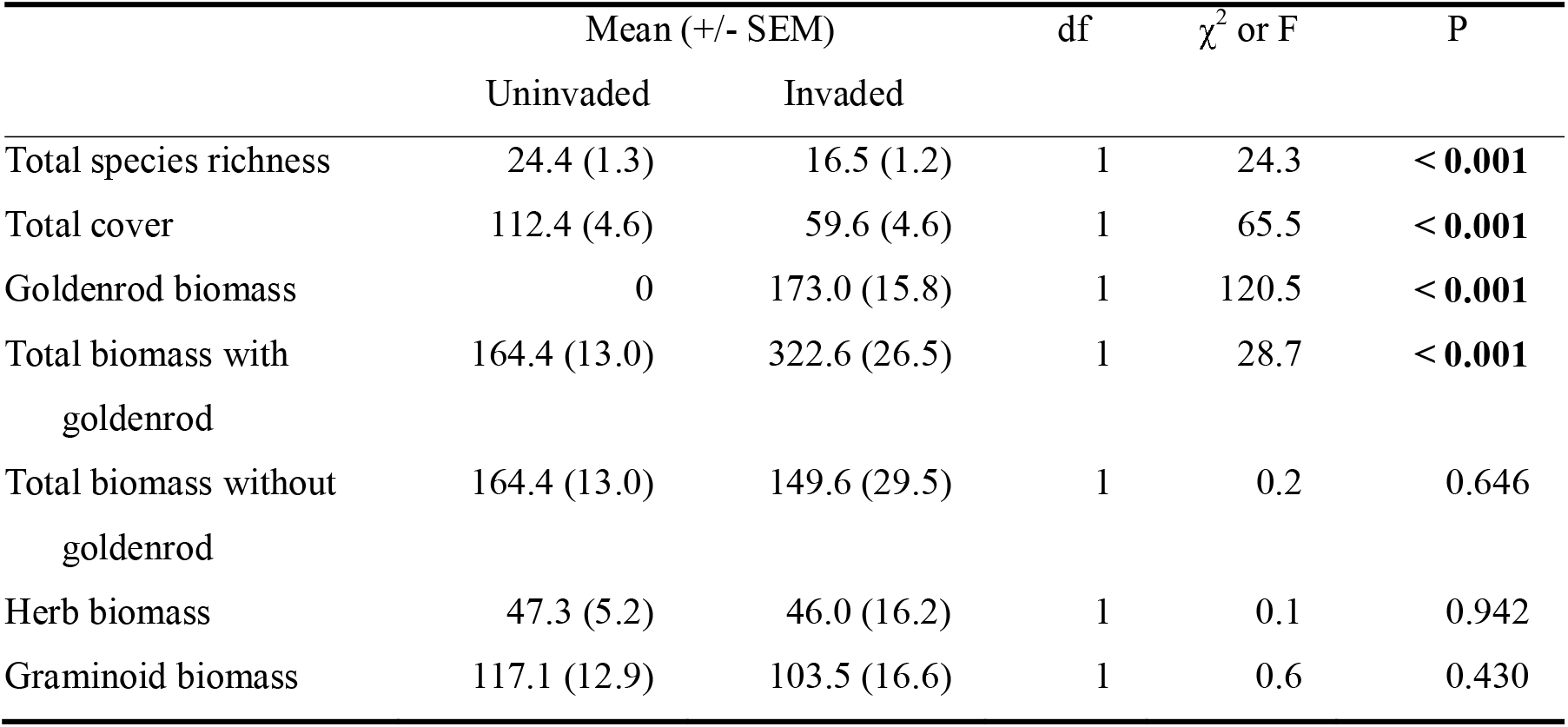
Group means of uninvaded and invaded plots with ANOVA table based on the linear mixed-effects models and generalized linear mixed-effects models for total species richness, total cover, and biomass data. Species richness, and cover data were analysed excluding goldenrod. Herb biomass included the minor (below 1%) biomass of phanerophyte species. P values less than 0.05 shown in bold.

According to our results, the average absolute total cover without goldenrod was also reduced by 47.0% compared to control plots. Despite the large difference in total cover, we did not find any significant effect on either total biomass without goldenrod, or herb or graminoid biomass. In a mesocosm experiment, the invaded community-level biomass (without goldenrod) was not significantly different from that in uninvaded vegetation (Scharfy et al., 2010). However, when analysing total biomass with goldenrod (i.e. community-level biomass), we can see a large (96.2%) increase in invaded plots. This can be attributed to the larger shoot height and high shoot density of the two goldenrod species (Ye et al., 2019). The high productivity caused by high nutrient (P and K) accumulation of the species may explain their large biomass (Ye et al., 2019). Biomass increase may also be explained by the fast inherent growth rate of goldenrods, their potential exploitation of unused resources, and a more efficient use of nutrients for biomass production (Scharfy et al., 2009).

Although goldenrod invasion increased overall primary productivity, it did not altered hay composition in terms of changing herb or graminoid biomass. Szymura et al. (2022) also showed that neither forb nor graminoid biomass changed following goldenrod control with herbicide application. As opposed to this, giant goldenrod was shown to reduce the quality of hay because of its high saponin content (Weber & Jakobs, 2005). Both goldenrod invasion and the improvement of hay quality can be increased by the application of fresh hay as a seed source in vegetation restoration of goldenrod invaded fields (Szymura et al., 2022).

As for origin of species, goldenrod invasion decreased only the richness but not the cover of native and exotic species from 23.1 to 16.3 and from 1.4 to 0.3, respectively (Table 2, Fig. 1). This large decrease may be attributed to the high competitive ability of goldenrods. However, it should be noted that the net effect of goldenrods on native species richness can range from negative to positive depending on the quality (species diversity and evenness) of the recipient community, the dominance of the invader, and also on the climate (Dong et al., 2015). Negative effects on native species may also be the results of allelopathic activity of the roots that were shown to be stronger on species in invaded range than in the native range (Pál et al., 2015).

**TABLE 2.** ANOVA table of the linear mixed-effects models and generalized linear mixed-effects models for species richness, cover, and biomass data. Species richness and species cover data were analysed excluding goldenrod. P values less than 0.05 are shown in bold.

|  | Species richness |  |  | Percentage cover |  |  |
| --- | --- | --- | --- | --- | --- | --- |
| | df | $\chi^2$ value | P | df | F value | P |
| *Origin* |  |  |  |  |  |  |
| Native | 1 | 18.7 | **< 0.001** | 1 | 3.1 | 0.069 |
| Exotic | 1 | 9.8 | **0.002** | 1 | 3.1 | 0.069 |
| *Life span* |  |  |  |  |  |  |
| Short-lived | 1 | 13.9 | **< 0.001** | 1 | 0.1 | 0.805 |
| Perennial | 1 | 13.9 | **< 0.001** | 1 | 0.1 | 0.855 |
| *Growth form* |  |  |  |  |  |  |
| Herb | 1 | 13.6 | **< 0.001** | 1 | 0.1 | 0.895 |
| Legume | 1 | 3.6 | 0.060 | 1 | 3.2 | 0.073 |
| Graminoid | 1 | 8.3 | **0.004** | 1 | 0.1 | 0.820 |
| Phaneropyte | 1 | 0.9 | 0.341 | 1 | 0.9 | 0.338 |
| *Social Behaviour Types* |  |  |  |  |  |  |
| Specialist | 1 | 2.2 | 0.142 | 1 | 2.4 | 0.118 |
| Generalist | 1 | 6.0 | **0.014** | 1 | 2.3 | 0.129 |
| Competitor | 1 | 16.5 | **< 0.001** | 1 | 1.4 | 0.229 |
| Ruderal | 1 | 8.6 | **0.003** | 1 | 5.9 | **0.015** |

**FIGURE 1.**
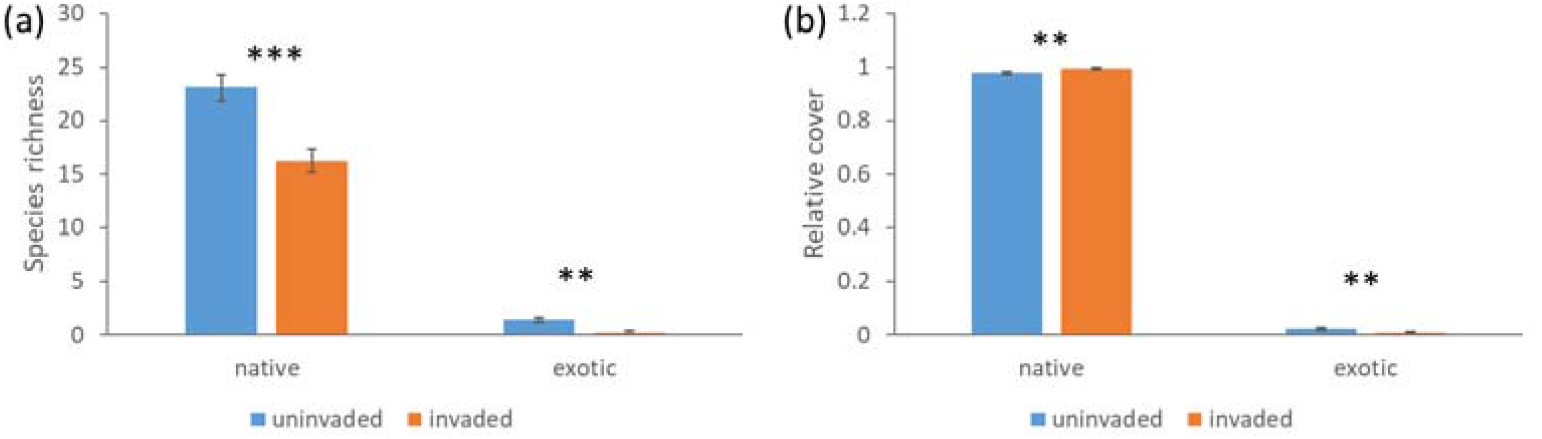
(a) Richness (Mean ± SEM) and (b) relative percentage cover (Mean ± SEM) of native and exotic species (apart from goldenrod) in uninvaded and invaded relevés (n=16).

Similarly, goldenrod presence impacted the richness of both short-lived and perennial species negatively, resulting in 52.6% and 27.7% decrease, respectively (Fig. 2). This effect was similar to the results of Bielecka et al. (2020) who found that the richness of both annuals and perennials decreased in the invaded sites in Poland. In contrast, the relative cover of the two life span groups was not affected by goldenrod invasion as most of the cover was made up by perennial species in both invaded and uninvaded plots.

**FIGURE 2.**
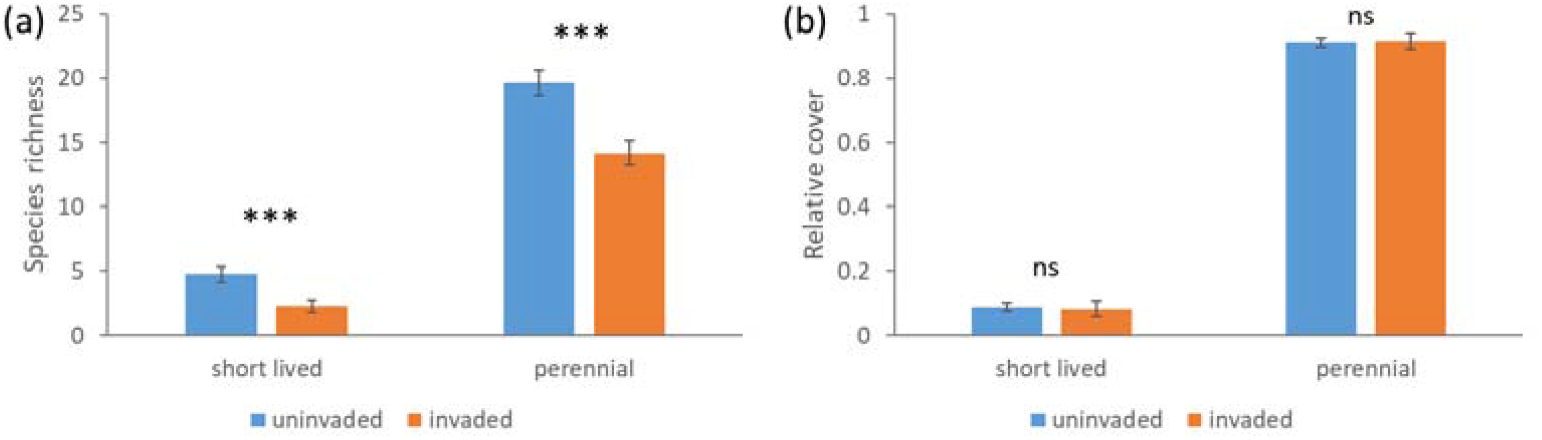
(a) Richness (Mean ± SEM) and (b) relative percentage cover (Mean ± SEM) of short-lived (annuals and biennials) and perennial species (apart from goldenrod) in uninvaded and invaded relevés (n=16).

Fig. 3 shows that life form distribution was affected only in terms of species richness but not in terms of relative cover. Herb and graminoid richness was significantly lower in goldenrod invaded plots, whereas legumes and phanerophytes did not show any differences in the two groups. Dong et al. (2015) discussed the possibility that goldenrods only displace dominant species without affecting non-dominant species, thus allowing for an unchanged species richness. In our case, however, we observed a similar suppressing effect of goldenrod invasion on both dominant graminoids and non-dominant herbs.

**FIGURE 3.**
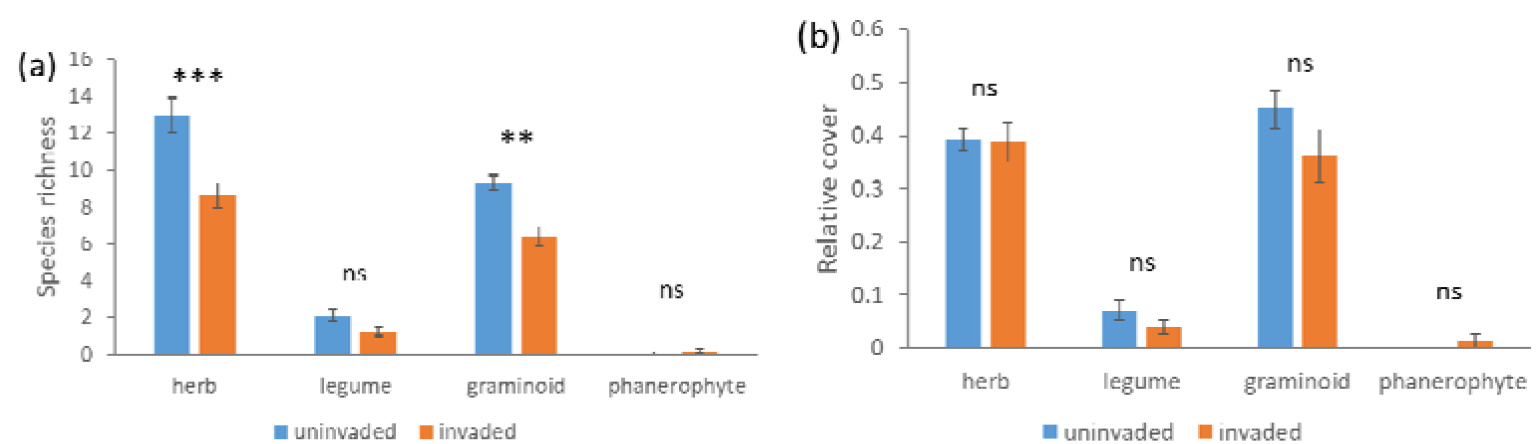
(a) Richness (Mean ± SEM) and (b) relative percentage cover (Mean ± SEM) of herbs (without legumes and goldenrod), legumes, graminoid (*Poaceae* and *Cyperaceae*), and phanerophyte species in uninvaded and invaded relevés (n=16).

The richness and cover of specialist species was similarly low in both invaded and uninvaded vegetation (Fig. 4). Similarly, Fenesi et al. (2015) found no effect of Canadian goldenrod cover on the diversity of grassland specialists. Furthermore, we found that the species richness of generalists, dominant competitors, and ruderals decreased following goldenrod invasion. In comparison, Fenesi et al. (2015) found no effect of Canadian goldenrod cover on the richness of ruderal species. In their study, old field age had a dominant effect over goldenrod cover related to this species group. In terms of generalist species diversity, however, they found similar pattern, as goldenrod displaced native generalist species in young successional old fields. In other case, despite the decrease of species richness of ruderals, we detected a significant increase in their cover in the invaded plots at the expense of the other three species groups. This showed a reduction in the naturalness and conservation value of the invaded stands.

**FIGURE 4.**
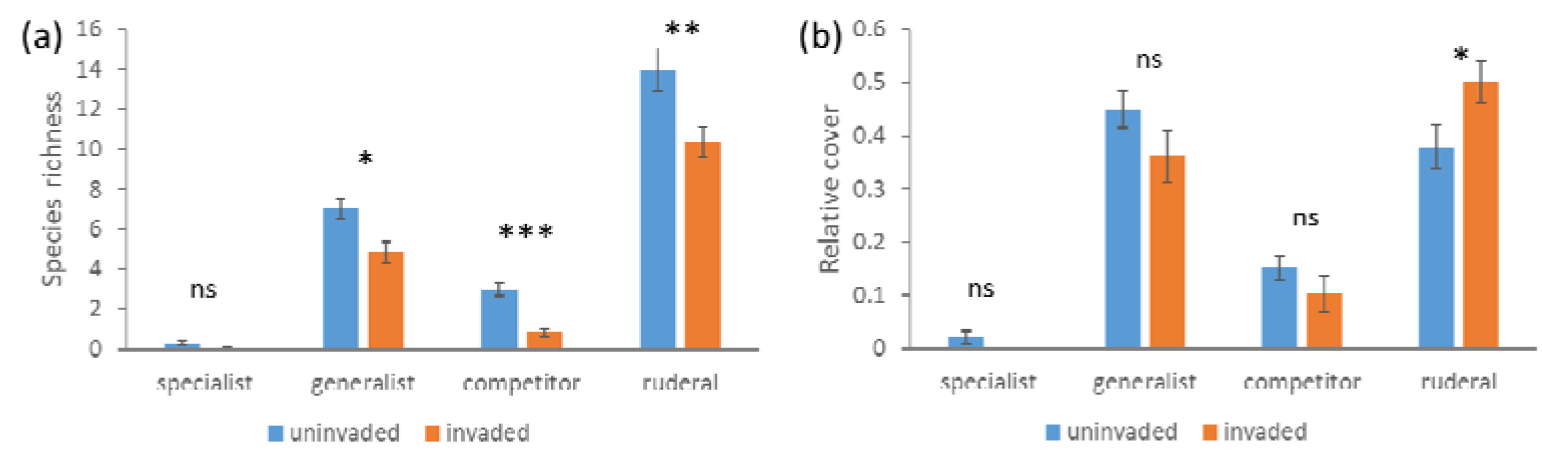
(a) Richness (Mean ± SEM) and (b) relative percentage cover (Mean ± SEM) of Social Behaviour Types (Borhidi, 1995) in uninvaded and invaded relevés. Ruderals include natural pioneers, disturbance tolerants, weeds, ruderal competitors, and alien species (n=16).

Our results showed that goldenrod invasion had significant effects on the species richness and total biomass of wet meadow vegetation. The found a decline in nature conservation value due to goldenrod invasion in terms of the shift in species composition from specialist, generalist and dominant species toward ruderal species. Furthermore, although the invasion resulted in the decrease of native species richness, it also affected other exotic species negatively. As for forage quantity, we found a significant increase in community-level biomass. However, this increase could be attributed solely to the biomass of the invader itself, as the biomass of the recipient species was similar in the invaded and uninvaded plots, and we did not observe a shift in biomass towards either herb or graminoid dominance.

## 5 CONCLUSIONS

In conclusion, the study revealed that goldenrod invasion had a negative impact on species richness and community-level biomass in wet meadows. The invasion led to a decrease in native and exotic species richness, a shift towards ruderal species, and an increase in total biomass attributed mainly to the invader itself.

## ACKNOWLEDGEMENTS

This study was supported by the Hungarian Széchenyi 2020 Programme (VP3-16.1.1-4.1.5 No. 3043361958, 1907562176), and the Hungarian National Research, Development and Innovation Office (NKFIH KKP133839).

## Notes

### Competing Interest Statement

The authors have declared no competing interest.

