## Supplementary material for "Effects of goldenrod (*Solidago gigantea* Aiton and S. *canadensis* L.) invasion on the wet meadow vegetation composition of Somogyfajsz wood pasture": Table S1

**Supplementary information**

Table S1. Analysed plant species with their analysed characteristics, and their frequency in invaded and uninvaded plots (n=16). Species were assigned to functional groups based on their origin (native vs exotic), life span (short-lived, or perennial), growth form (graminoid, legume, herb, or phanerophyte), and Social Behaviour Types (SBT). Abbreviations: C: competitor, S: specialist, G: generalist, R: ruderal. NA denotes missing values.

| Species number | Origin | Lifespan | Groth form | SBT | Invaded | Uninvaded |
| --- | --- | --- | --- | --- | --- | --- |
| *Achillea sp.* | native | perennial | herb | ruderal | 7 | 9 |
| *Agrostis stolonifera* | native | perennial | graminoid | competitor | 1 | 7 |
| *Allium scorodoprasum* | native | perennial | herb | ruderal | 1 | 0 |
| *Alnus glutinosa* | native | perennial | phanerophyte | competitor | 1 | 0 |
| *Ambrosia artemisiifolia* | exotic | short-lived | herb | ruderal | 1 | 7 |
| *Anacamptis morio* | native | perennial | herb | generalist | 0 | 1 |
| *Anthoxanthum odoratum* | native | perennial | graminoid | competitor | 1 | 9 |
| *Aster linosyris* | native | perennial | herb | generalist | 2 | 2 |
| *Briza media* | native | perennial | graminoid | generalist | 4 | 6 |
| *Bromus hordeaceus* | native | short-lived | graminoid | ruderal | 4 | 2 |
| *Calamagrostis epigeios* | native | perennial | graminoid | ruderal | 2 | 0 |
| *Carex distans* | native | perennial | graminoid | competitor | 4 | 10 |
| *Carex flacca* | native | perennial | graminoid | generalist | 0 | 1 |
| *Carex fritschii* | native | perennial | graminoid | specialist | 0 | 1 |
| *Carex hirta* | native | perennial | graminoid | ruderal | 14 | 13 |
| *Carex panicea* | native | perennial | graminoid | generalist | 6 | 7 |
| *Carex spicata* | native | perennial | graminoid | ruderal | 5 | 4 |
| *Caryophyllaceae* | native | NA | herb | NA | 1 | 0 |
| *Caryophyllaceae* | native | NA | herb | NA | 0 | 1 |
| *Centaurea jacea* | native | perennial | herb | generalist | 4 | 14 |
| *Cerastium sp.* | native | short-lived | herb | ruderal | 0 | 10 |
| *Cirsium arvense* | native | perennial | herb | ruderal | 2 | 1 |
| *Convolvulus arvensis* | native | perennial | herb | ruderal | 0 | 1 |
| *Crataegus monogyna* | native | perennial | phanerophyte | generalist | 1 | 0 |
| *Cruciata pedemontana* | native | short-lived | herb | generalist | 6 | 1 |
| *Cynodon dactylon* | native | perennial | graminoid | ruderal | 1 | 6 |
| *Dactylis glomerata* | native | perennial | graminoid | ruderal | 10 | 6 |
| *Danthonia decumbens* | native | perennial | graminoid | specialist | 0 | 4 |
| *Daucus carota* | native | short-lived | herb | ruderal | 5 | 14 |
| *Deschampsia caespitosa* | native | perennial | graminoid | competitor | 1 | 1 |
| *Equisetum arvense* | native | perennial | herb | ruderal | 3 | 3 |
| *Erigeron annuus* | exotic | short-lived | herb | ruderal | 3 | 13 |
| *Festuca pratensis* | native | perennial | graminoid | competitor | 2 | 12 |
| *Festuca rubra* | native | perennial | graminoid | competitor | 3 | 7 |
| *Frangula alnus* | native | perennial | phanerophyte | generalist | 0 | 1 |
| *Fraxinus pennsylvanica* | exotic | perennial | herb | ruderal | 0 | 2 |
| *Galium mollugo* | native | perennial | herb | generalist | 2 | 0 |
| *Galium verum* | native | perennial | herb | ruderal | 15 | 12 |
| *Glechoma hederacea* | native | perennial | herb | ruderal | 6 | 6 |
| *Helictotrichon pubescens* | native | perennial | graminoid | generalist | 8 | 12 |
| *Hieracium sp.* | native | perennial | herb | NA | 0 | 1 |
| *Holcus lanatus* | native | perennial | graminoid | generalist | 13 | 16 |
| *Hypericum perforatum* | native | perennial | herb | ruderal | 0 | 3 |
| *Juncus sp.* | native | perennial | graminoid | NA | 1 | 0 |
| *Koeleria cristata* | native | perennial | graminoid | generalist | 0 | 1 |
| *Lotus corniculatus* | native | perennial | legume | ruderal | 1 | 9 |
| *Luzula campestris* | native | perennial | graminoid | ruderal | 5 | 11 |
| *Lycopus exaltatus* | native | perennial | herb | ruderal | 1 | 0 |
| *Medicago minima* | native | short-lived | legume | generalist | 0 | 7 |
| *Mentha aquatica* | native | perennial | herb | generalist | 3 | 3 |
| *Molinia caerulea* | native | perennial | graminoid | competitor | 1 | 2 |
| *Myosotis arvensis* | native | short-lived | herb | ruderal | 1 | 0 |
| *Ophioglossum vulgatum* | native | perennial | herb | generalist | 1 | 0 |
| *Origanum vulgare* | native | perennial | herb | ruderal | 8 | 6 |
| *Peucedanum arenarium* | native | perennial | herb | specialist | 1 | 0 |
| *Peucedanum oreoselinum* | native | perennial | herb | generalist | 1 | 3 |
| *Plantago lanceolata* | native | perennial | herb | ruderal | 3 | 11 |
| *Poa pratensis* | native | perennial | graminoid | generalist | 12 | 11 |
| *Poa trivialis* | native | perennial | graminoid | ruderal | 2 | 0 |
| *Poaceae* | native | perennial | graminoid | NA | 3 | 0 |
| *Polygala comosa* | native | perennial | herb | generalist | 1 | 1 |
| *Potentilla heptaphylla* | native | perennial | herb | generalist | 0 | 1 |
| *Potentilla reptans* | native | perennial | herb | ruderal | 10 | 8 |
| *Prunella vulgaris* | native | perennial | herb | ruderal | 1 | 5 |
| *Ranunculus acris* | native | perennial | herb | generalist | 5 | 7 |
| *Ranunculus repens* | native | perennial | herb | ruderal | 6 | 8 |
| *Rhinanthus serotinus* | native | short-lived | herb | generalist | 8 | 14 |
| *Rosa canina* | native | perennial | phanerophyte | ruderal | 1 | 0 |
| *Rumex acetosella* | native | perennial | herb | ruderal | 11 | 9 |
| *Saxifraga bulbifera* | native | perennial | herb | generalist | 1 | 2 |
| *Solidago gigantea* | exotic | perennial | herb | ruderal | 16 | 0 |
| *Stellaria graminea* | native | perennial | herb | ruderal | 0 | 2 |
| *Taraxacum officinale* | native | perennial | herb | ruderal | 6 | 8 |
| *Trifolium pratense* | native | perennial | legume | ruderal | 5 | 14 |
| *Trifolium repens* | native | perennial | legume | ruderal | 1 | 0 |
| *Verbascum nigrum* | native | short-lived | herb | ruderal | 0 | 1 |
| *Verbena officinalis* | native | short-lived | herb | ruderal | 0 | 3 |
| *Veronica chamaedrys* | native | perennial | herb | ruderal | 12 | 10 |
| *Veronica serpyllifolia* | native | perennial | herb | ruderal | 0 | 1 |
| *Veronica sp.* | native | short-lived | herb | ruderal | 0 | 2 |
| *Vicia angustifolia* | native | short-lived | legume | ruderal | 3 | 0 |
| *Vicia cracca* | native | perennial | legume | ruderal | 5 | 2 |
| *Vicia grandiflora* | native | short-lived | legume | ruderal | 2 | 1 |
| *Vicia hirsuta* | native | short-lived | legume | ruderal | 3 | 1 |
| *Viola sp.* | native | perennial | herb | generalist | 0 | 1 |
